# Design of universal Ebola virus vaccine candidates *via* immunofocusing

**DOI:** 10.1101/2023.10.14.562364

**Authors:** Duo Xu, Abigail E. Powell, Ashley Utz, Mrinmoy Sanyal, Jonathan Do, J.J. Patten, Juan I. Moliva, Nancy J. Sullivan, Robert A. Davey, Peter S. Kim

## Abstract

Ebola virus causes hemorrhagic fever in humans and poses a significant threat to global public health. Although two viral vector vaccines have been approved to prevent Ebola virus disease, they are distributed in the limited ring vaccination setting and only indicated for prevention of infection from *orthoebolavirus zairense* (EBOV) – one of three *orthoebolavirus* species that have caused previous outbreaks. Ebola virus glycoprotein GP mediates viral infection and serves as the primary target of neutralizing antibodies. Here we describe a universal Ebola virus vaccine approach using structure-guided design of candidates with hyperglycosylation that aims to direct antibody responses away from variable regions and toward conserved epitopes of GP. We first determined the hyperglycosylation landscape on Ebola virus GP and used that to generate hyperglycosylated GP variants with two to four additional glycosylation sites to mask the highly variable glycan cap region. We then created vaccine candidates by displaying wild-type or hyperglycosylated GP variants on ferritin nanoparticles (Fer). Immunization with these antigens elicited potent neutralizing antisera against EBOV in mice. Importantly, we observed consistent cross-neutralizing activity against Bundibugyo virus and Sudan virus from hyperglycosylated GP-Fer with two or three additional glycans. In comparison, elicitation of cross-neutralizing antisera was rare in mice immunized with wild-type GP-Fer. These results demonstrate a potential strategy to develop universal Ebola virus vaccines that confer cross-protective immunity against existing and emerging filovirus species.

**SIGNIFICANCE STATEMENT:** Ebola virus outbreaks cause hemorrhagic fever with high mortality rates. Current viral vaccines require cold-chain storage and are distributed in limited ring vaccination settings. They are only indicated for protection against *orthoebolavirus zairense* (EBOV), one of three human-pathogenic Ebola virus species. Here we harness hyperglycosylation as an immunofocusing approach to design universal Ebola virus vaccine candidates based on Ebola virus glycoprotein (GP) displayed on ferritin nanoparticles (Fer). Compared with wild-type GP-Fer, immunization with hyperglycosylated GP-Fer elicited potently neutralizing antisera against EBOV, and more importantly, consistent cross-neutralizing activity against the other two orthoebolavirus species. Our work shows that immunofocusing antibody responses toward conserved and neutralizing epitopes of GP represents a promising strategy for vaccine design against antigenically diverse Ebola virus species.

## INTRODUCTION

Ebola virus, a member of the *Filoviridae* family, is highly pathogenic and can cause hemorrhagic fever in humans with severe morbidity and high mortality(1). Since its discovery in 1976, Ebola virus has caused more than 20 outbreaks in Africa, most notably the 2014-2016 epidemic that quickly became an international public health emergency(1–3). Two viral vector vaccines (Ervebo^®^ and Zabdeno/Mvabea^®^) have been approved for prevention of Ebola virus disease(4). However, neither vaccine is widely distributed for outbreak prevention since they both require cold-chain storage(5) and may cause mild to moderate side effects in vaccinated individuals(6, 7). Instead, they have only been used in limited “ring vaccination” settings to protect high-risk groups throughout endemic areas during active outbreaks(8, 9). Moreover, these two vaccines are only indicated for prevention of infection from *orthoebolavirus zairense* (EBOV)(10, 11), whereas three species of the *orthoebolavirus* genus have caused outbreaks and remain ongoing threats: EBOV, *orthoebolavirus bundibugyoense* (BDBV) and *orthoebolavirus sudanense* (SUDV)(2, 3). The existence of these antigenically different *orthoebolavirus* species necessitates the design of new prophylactic vaccines that are suitable for widespread use and confer durable and cross-protective immunity.

The trimeric EBOV glycoprotein (GP) is the sole viral surface protein and it mediates viral infection of host cells(12). Following viral uptake *via* endocytosis, GP is proteolytically processed in endosomes(13, 14), where the mucin-like domain and the glycan cap – two poorly conserved regions(15) – are cleaved to expose its receptor-binding domain, allowing its binding to the intracellular receptor Niemann-Pick C1(16, 17) (**Fig. 1a**). Subsequently, GP undergoes structural rearrangement to prompt fusion of viral and cellular membranes and transfer of the viral genome into the cytosol(18). Monoclonal antibodies (mAbs) that bind GP and block viral entry have been shown to prevent EBOV infection in nonhuman primates (NHPs)(19–21) and humans(22, 23), with two antibody drugs (Ebanga^®^ and Inmazeb^®^) approved to treat Ebola virus disease(24, 25). Although these clinically approved mAbs are only indicated for EBOV, several other GP-targeting mAbs have been isolated that can neutralize all three *orthoebolavirus* species(26–29), suggesting that conserved and cross-reactive epitopes exist on GP.

**Figure 1.**
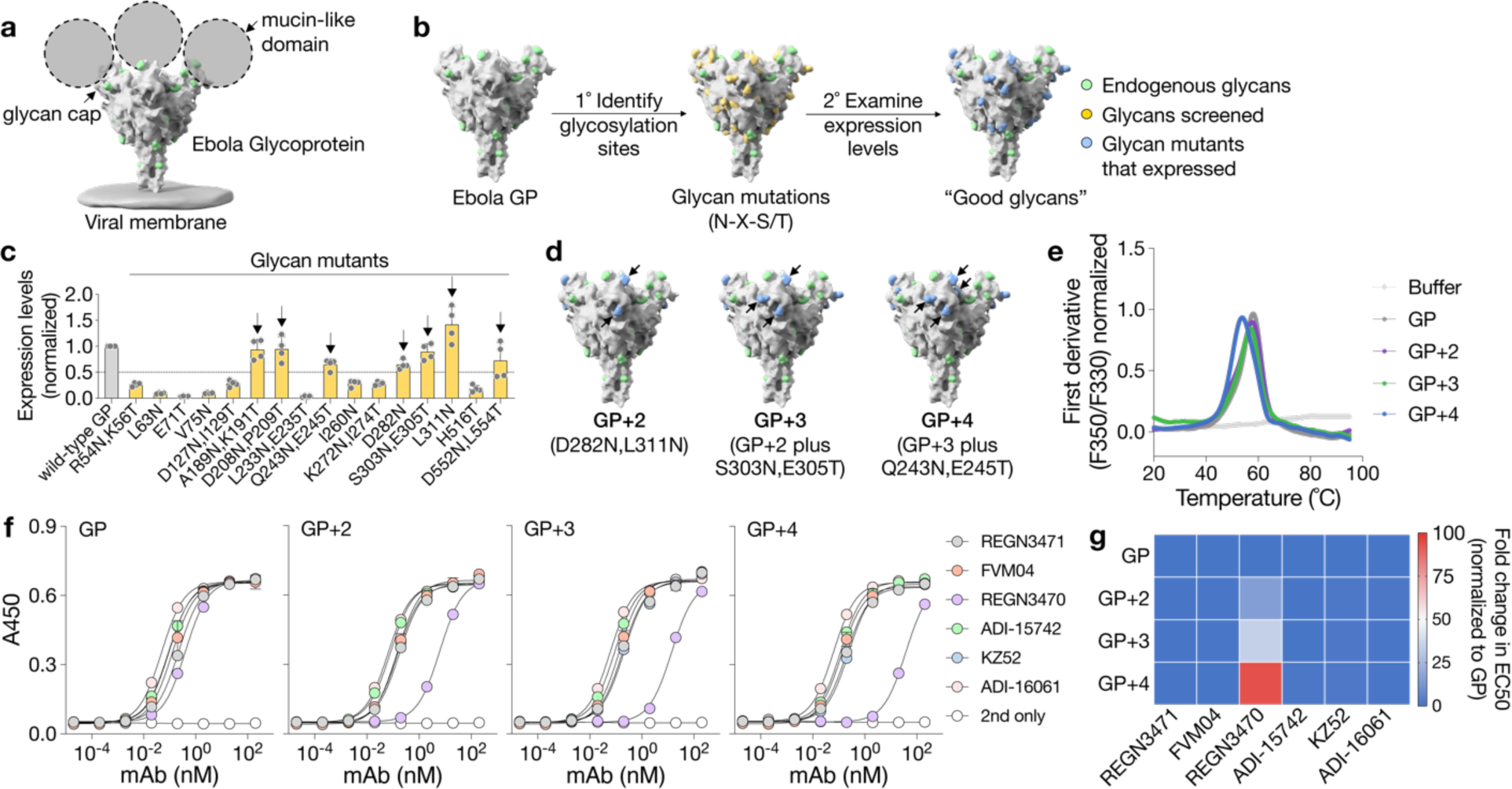
Glycosylation landscape on EBOV GP. **a**, Structure of the trimeric EBOV GP prior to endosomal processing (PDB ID 5JQ3, grey). Mucin-like domain (dashed line) and glycan cap regions are proteolytically cleaved in endosomes during viral entry into host cells. The ectodomain of EBOV GP with the mucin-like domain deleted is referred as EBOV GP. Green spheres indicate endogenous glycans. **b**, Screening and identification of permissive glycan installations on EBOV GP trimers (PDB ID 5JQ3, grey). Individual *N*-linked glycosylation sites with Asn-X-Ser/Thr (N-X-S/T, where X can be any amino acid except proline) motifs were introduced by site-directed mutagenesis and indicated by yellow spheres on each GP protomer. Introduction of N-X-S/T motifs involved mutating one or two residues in the sequence. Single-glycan mutants were then transiently expressed in Expi-293F cells, and their expression levels were analyzed by Western-blotting. **c**, Expression levels of single-glycan mutants were analyzed by Western blots and normalized to wild-type GP. Arrows indicate “good glycans” above the dashed line, defined as >50% of wild-type GP expression. Data are presented as mean ± standard deviation (*n*=4 biological replicates). **d**, Hyperglycosylated GP with two, three or four *N*-linked glycans (blue spheres indicated by arrows) installed in the glycan cap region of each GP protomer. **e**, Representative thermal melting profiles of GP and its glycan variants measured by differential scanning fluorimetry. **f**, Epitope analysis of GP and its glycan variants with six GP-specific mAbs by ELISA. Binding data are presented as mean ± standard deviation (*n*=2 biological replicates with technical duplicates). **g**, Fold-change in binding (EC_50_) of GP-specific mAbs (columns) to GP and its glycan variants (rows). EC_50_ (antibody concentration with half-maximal binding) was calculated from (**g)** and normalized to values obtained from the wild-type GP group.

Although natural infection and vaccination with GP-based immunogens have been shown to elicit neutralizing antibody responses in animals(30–37) and humans(38, 39), they only induce limited cross-reactivity among different *orthoebolavirus* species(40). Therefore, a direct approach to achieve cross-protection is to combine individual vaccines(41–44). For instance, a heterologous prime-boost regimen with recombinant vesicular stomatitis virus (rVSV)-based vaccines expressing EBOV GP and SUDV GP protected against BDBV challenge in NHPs, while a single-dose blended vaccination failed to provide protection(42), suggesting that immune responses to a bivalent vaccine may bias toward certain epitopes over others(40). Compared with multivalent viral vaccines, we envision that a single protein-based antigen that protects against all three *orthoebolavirus* species that have caused previous outbreaks represents a more attractive alternative, which would be easier to manufacture and store than viral vaccines and could enable widespread vaccination strategies beyond ring vaccination.

EBOV vaccine candidates that focus antibody responses toward conserved epitopes on GP are likely to elicit antisera with broad binding activity or even cross-neutralization. Such candidates may serve as the basis of a universal Ebola virus vaccine. Here we harness hyperglycosylation as an immunofocusing approach for vaccine design(45–48), which acts by masking variable or non-neutralizing epitopes (*i.e.*, the glycan cap) by installation of glycans, thereby re-directing antibody responses to conserved or neutralizing epitopes. Based on the structure of EBOV GP with the mucin-domain deleted (referred as EBOV GP), we first screened and identified permissive sites for hyperglycosylation. We then installed different numbers (two, three, or four) of additional glycosylation sites on each GP protomer to mask the poorly conserved glycan cap region, and generated three hyperglycosylated GP variants (GP+2, GP+3 and GP+4). We displayed these GP variants on ferritin nanoparticles (GP-Fer), which have been previously shown to dramatically enhance the immunogenicity of protein-based antigens, including for influenza A virus(49, 50), human immunodeficiency virus-1 (HIV-1)(51, 52), and severe acute respiratory syndrome coronavirus-2 (SARS-CoV-2)(53, 54). While wild-type and hyperglycosylated GP-Fer induced neutralizing responses against EBOV in immunized mice, only GP+2-Fer and GP+3-Fer consistently delivered superior cross-neutralizing activity against BDBV and SUDV, forming the basis of new universal Ebola virus vaccine candidates.

## RESULTS

### Glycosylation landscape on EBOV GP

Based on the X-ray crystal structure of EBOV GP(55), we selected 16 potential sites that are exposed on the protein surface for prospective glycan installations, and then created individual glycan mutants with inferred *N-*linked glycosylation sites *via* site-directed mutagenesis (**Fig. 1b**). We analyzed the expression level of those single-glycan variants by Western blots (**Fig. S1a**) and compared them with wild-type GP to identify “good glycans” whose installation did not lead to significant protein misfolding or non-expression. While more than half of the mutations greatly reduced GP expression in Expi293F cells (**Fig. 1c**), seven mutants maintained >50% of GP expression compared with the wild-type. Four of these “good glycans” were located around the glycan cap region, with the other three toward the base of the GP trimer (**Fig. S1b**). Based on the sequence conservation of GP among different *orthoebolavirus* species (**Fig. S1c**), we preferentially installed glycans around the glycan cap to mask this poorly conserved region. By combining single-glycan mutants, we installed two, three or four glycans on each GP protomer (GP+2, GP+3 or GP+4) to afford different levels of epitope masking (**Fig. 1d**). These glycan variants were successfully expressed as trimers and purified to homogeneity *via* affinity purification followed by gel filtration (**Fig. S1d**). Compared to wild-type GP, they exhibited slower migration patterns on gel electrophoresis and Western blotting, indicating the successful installation of additional glycans (**Fig. S1e**). GP, GP+2 and GP+3 also shared similar thermal melting profiles (**Fig. 1e**) with melting temperatures (T_m_, **Fig. S1f**) around 58°C. The T_m_ of GP+4 was lower (54 °C), suggesting that installation of four glycans may slightly destabilize the GP structure, although all proteins were thermostable at physiological temperature. To examine the degree of epitope-masking, we measured binding of six GP-specific mAbs to wild-type and hyperglycosylated GPs with enzyme-linked immunosorbent assays (ELISAs) (**Fig. 1f**). These mAbs recognize unique epitopes on GP (REGN3471 – head(20), FVM04 – head(56), REGN3470 – glycan cap(20), ADI-15742 – fusion loop(27), KZ52 – GP base(15), ADI-16061 – heptad repeat-2(27)) and allow for rapid epitope-mapping (**Fig. S1g**). We observed no noticeable difference in mAb-binding to GP and its variants, except for REGN3470 that targets the glycan cap(20) (**Fig. 1g**). As we increased the number of glycans, binding of REGN3470 toward GP variants gradually decreased, approaching a 93-fold difference in EC_50_ (mAb concentration with half-maximal binding) for GP+4 compared to wild-type GP. Having established the glycosylation landscape on GP and created hyperglycosylated GP variants with different levels of epitope-masking of the glycan cap, we set out to generate nanoparticle-based vaccine candidates for the subsequent assessment of immunogenicity.

### Display of GP on ferritin nanoparticles

Although protein-based subunit vaccines are often less immunogenic than their viral vector counterparts, multivalent presentation of antigens on nanoparticles dramatically enhances antigen valency and immunogenicity(57). We therefore displayed GP and its glycan variants on *Helicobacter pylori* ferritin nanoparticles (Fer), which presented eight copies of the GP trimer on the surface at the threefold axes of symmetry(58) (Fig 2a). GP-Fer and its glycan variants (GP+2-Fer, GP+3-Fer, and GP+4-Fer) were expressed and purified to homogeneity *via* anion-exchange chromatography followed by gel filtration (**Fig. S2a**). We found that glycan installations increased the molecular weight of GP-Fer, as measured by gel electrophoresis and Western blotting (**Fig. S2b**). Display of GP and its variants on Fer did not significantly affect their thermal melting profiles (**Fig. S2c**) or T_m_ (Fig. 2b), probably because Fer unfolds at a high temperature (∼80°C, **Fig. S2c**). Similar to GP+4 (Fig. 1d), the T_m_ of GP+4-Fer was also lower than that of GP-Fer, GP+2-Fer or GP+3-Fer. GP-Fer and its glycan variants (GP+2-Fer and GP+3-Fer) also maintained their thermal stability at 37°C for up to 14 days (**Fig. S2d,e**). In addition, consistent with the previous ELISA results (Fig. 1e, f), REGN3470 was the only mAb that gradually lost its GP-binding capacity as the number of glycans increased (Fig. 2c**, Fig. S2f**), while the other five mAbs showed similar binding profiles to GP-Fer and its glycan variants.

**Figure 2.**
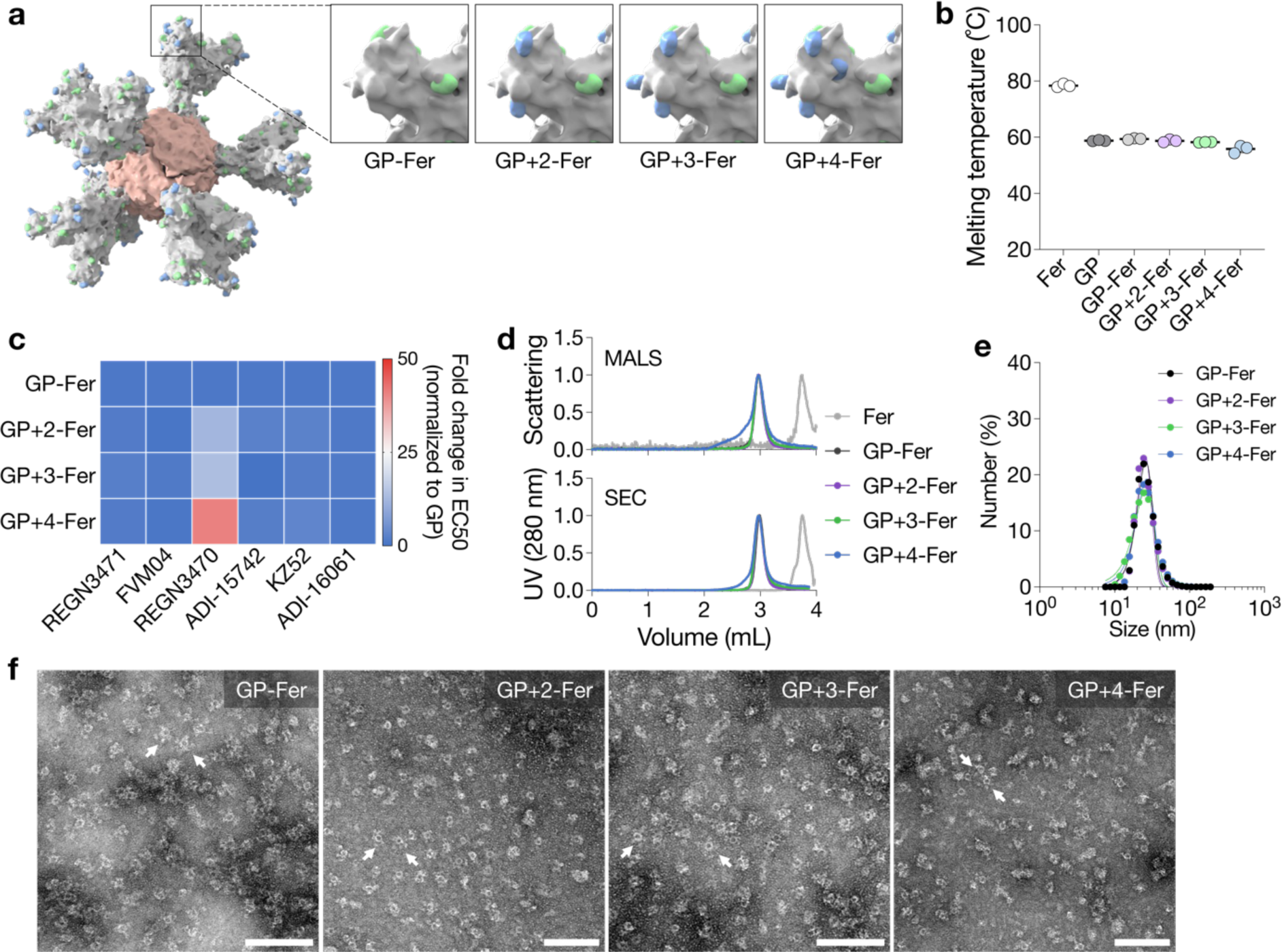
Display of GP on ferritin nanoparticles. **a**, Schematic illustration of eight copies of GP trimers displayed on one ferritin nanoparticle (Fer, PDB ID, 3BVE). Glycan installations on the glycan cap region were highlighted for GP-Fer and its glycan variants. Green spheres indicate endogenous glycans, and blue spheres indicate installation sites of additional glycans. **b**, Thermal melting temperatures of wild-type Fer, GP-Fer and its glycan variants measured by differential scanning fluorimetry. Data are presented as mean ± standard deviation (*n*=3 biological replicates). **c**, Epitope analysis of GP-Fer and its glycan variants (rows) with six GP-specific mAbs (columns). Fold-change in binding (EC_50_) of GP-specific mAbs (columns) to GP-Fer and its glycan variants (rows). EC_50_ was calculated from Fig. S2f and normalized to values obtained from the wild-type GP-Fer group. **d**, Homogeneity of purified Fer, GP-Fer and its glycan variants analyzed by multi-angle light scattering coupled with size-exclusion chromatography (SEC-MALS). **e**, Size distribution GP-Fer and its glycan variants measured by dynamic light scattering. **f**, Representative transmission electron microscopy images of GP-Fer and its glycan variants. Arrows indicate representative protein nanoparticles. Scale bars, 100 nm.

We next examined the homogeneity of GP-Fer and its glycan variants with size exclusion chromatography coupled with multi-angle light scattering (SEC-MALS) using wild-type Fer as a control. All samples eluted as single peaks indicative of monodisperse nanoparticles (Fig. 2d). Elution profiles and refractive indices from MALS also informed the molecular weight (Mw) of GP-Fer and its glycan variants (**Fig. S2g,h**), where successive installation of glycans increased the Mw from 1.7 (GP-Fer) to 2.0 (GP+4-Fer) megadaltons. Consistent with MALS, dynamic light scattering confirmed the monodispersity of these protein nanoparticles with an average hydrodynamic diameter of 22 nm (Fig. 2e**, Fig. S2i**). Finally, we visualized the display of GP trimers on self-assembled Fer under transmission electron microscopy (Fig. 2f**, Fig. S2j**). Taken together, these results confirm the successful installation of GP and its glycan variants on Fer nanoparticles and demonstrate that glycan installations maintain the thermostability and antigenicity of GP.

### Immunization with GP-Fer and its glycan variants

To investigate whether multimerization on Fer increased immunogenicity, we immunized two groups of mice with GP trimer or GP-Fer adjuvanted with monophosphoryl lipid A(59) and Quil-A(60) (MPLA/Quil-A, Fig. 3a). A prime-boost regimen with GP-Fer elicited a rapid response (**Fig. S3a**) in mice and significantly higher EBOV GP-specific IgG titers than did the GP trimer three weeks post-boost (Fig. 3b). To determine whether increased antibody titers correlated with better neutralizing activity, we generated EBOV GP-pseudotyped lentiviruses (**Fig. S3b**) and performed neutralization assays with mouse antisera. Although both antigens induced neutralizing antisera against EBOV after the boost, the average neutralization titers (NT_50_) from the GP-Fer group were tenfold higher than the GP trimer group (Fig. 3c**, Fig. S3c,d**). These results are consistent with previous observations(49, 51, 61), where multimerization of antigens on nanoparticles enhanced neutralizing antibody responses.

**Figure 3.**
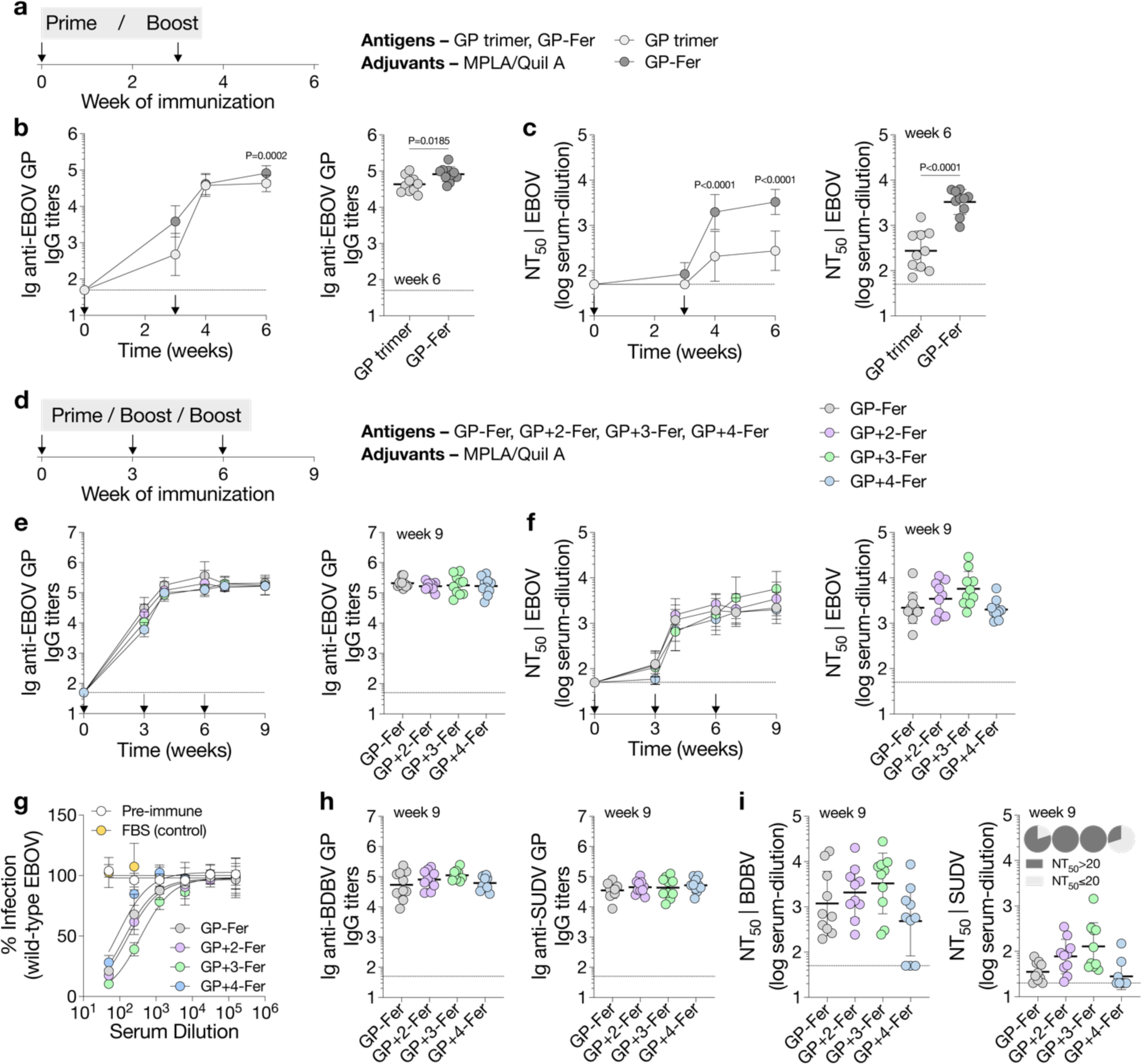
Immunogenicity of GP-Fer and its glycan variants in mice. **a**, Mouse immunization with a prime-boost regimen on days 0 and 21. Mice were immunized with 5 µg of antigens (GP trimer or GP-Fer) adjuvanted with MPLA/Quil-A (10 µg/10 µg) *via* subcutaneous injections (*n*=10 per group). **b**, EBOV GP-specific IgG titers over time (left). EBOV GP-specific IgG titers of individual mice on day 42 (week 6, right). **c**, Neutralization titers (NT_50_ – the serum dilution required to neutralize 50% of GP-pseudotyped lentiviruses) against EBOV over time (left). NT_50_ against EBOV from individual mice on day 42 (right). **d**, Mouse immunization with GP-Fer and its glycan variants (GP+2-Fer, GP+3-Fer or GP+4-Fer) with a three-dose regimen on days 0, 21 and 42. Mice were immunized with 5 µg of antigens adjuvanted with MPLA/Quil-A (10 µg/10 µg) *via* subcutaneous injections (*n*=10 per group). **e**, EBOV GP-specific IgG titers over time (left). EBOV GP-specific IgG titers of individual mice on day 63 (week 9, right). **f**, NT_50_ against EBOV over time (left). NT_50_ against EBOV from individual mice on day 63 (right). **g**, Neutralization of wild-type EBOV Mayinga with pooled mouse antisera from day 63 in each group (*n*=3 technical replicates). Pre-immune and fetal bovine serum (FBS) served as negative controls. **h**, BDBV GP-(left) or SUDV GP-specific (right) IgG titers of individual mice on day 63. **i**, NT_50_ against BDBV (left) or SUDV (right) from individual mice on day 63. Pie charts indicate the percentage of mice that developed neutralizing activity (NT_50_ over 20-fold serum dilution) against SUDV. Each circle represents a single mouse. Horizontal dashed lines indicate the limits of quantification. Data are presented as geometric mean ± standard deviation of log-transformed data in **b**,**c**,**e**,**f**,**h**,**i**. Comparisons of IgG titers or NT_50_ over time were performed using a two-way ANOVA followed by a Bonferroni test in **b,c**. Comparisons of two groups were performed using the two-tailed Mann-Whitney U test in **b,c**. *P* values of 0.05 or less are considered significant and plotted.

We then immunized four groups of mice with GP-Fer or its glycan variants (GP+2-Fer, GP+3-Fer or GP+4-Fer) adjuvanted with MPLA/Quil-A to investigate whether hyperglycosylated antigens could stimulate a cross-reactive response (Fig. 3d). Three weeks after the prime, mice from all groups developed EBOV GP-specific IgG responses, while titers in GP+3-Fer and GP+4-Fer groups were significantly lower than those in the GP-Fer group (**Fig. S4a**), indicating reduced immunogenicity resulting from hyperglycosylation. To determine whether the installed glycans masked antibody responses away from the glycan cap, we performed competition ELISAs, in which GP-coated ELISA plates were pre-incubated with a saturating concentration of glycan cap-targeting REGN3470 before adding antisera. While competition with REGN3470 led to a twofold decrease in binding titers of the GP-Fer group, we observed less competition as the number of glycans increased (**Fig. S4b**), suggesting that hyperglycosylation drove antibody responses away from the glycan cap – consistent with our design. The difference in antibody responses also correlated with NT_50_, where GP+4-Fer induced the lowest neutralizing activity among all antigens (**Fig. S4c**). Nonetheless, two booster injections of each antigen elevated antibody responses to high titers (Fig. 3e) with potent neutralizing activity against EBOV in all groups (Fig. 3f**, Fig. S4d**). Importantly, we further validated neutralization of EBOV using wild-type virus and observed similar potency with pooled antisera from each group (Fig. 3g).

We next analyzed binding of endpoint antisera (week 9) to BDBV and SUDV GP, and we found that all antigens induced cross-reactive antibodies to these two GPs (Fig. 3h) with titers about tenfold lower than those against EBOV GP. We also found that cross-reactive antisera competed with FVM04, a known cross-neutralizing mAb targeting GP-head(59) (**Fig. S4e**), in their binding to SUDV GP, while we observed less competition with ADI-15742, another cross-neutralizing mAb targeting the fusion loop(30). Using BDBV GP-or SUDV GP-pseudotyped lentiviruses (**Fig. S4f**), we then tested whether cross-reactive binding translated to cross-neutralization. All mice but three of ten in the GP+4-Fer group cross-neutralized BDBV (Fig. 3i**, Fig. S4g**). The average NT_50_ in GP+2-Fer and GP+3-Fer groups were also higher than those in GP-Fer and GP+4-Fer groups. We observed a similar trend in the cross-neutralization of SUDV (**Fig. S4h**) with GP+2-Fer and GP+3-Fer again outperforming GP-Fer and GP+4-Fer, although the neutralizing activity was modest. These results suggested that immunofocusing antibody responses away from the variable glycan cap and toward conserved epitopes afforded cross-neutralizing activity.

### Different immunization regimens with GP-Fer and its glycan variants

Although installation of four glycans afforded the best epitope masking of the glycan cap in mAb binding analysis (Fig. 1g), GP+4-Fer induced weaker cross-neutralizing activity than GP+2-Fer or GP+3-Fer did, suggesting that installation of the fourth glycan did not further boost the elicitation of cross-neutralizing antibodies, potentially due to over-masking of nearby epitopes or instability of GP+4-Fer as demonstrated by a decrease in its T_m_ (Fig. 2b). We thus continued our investigation of different immunization regimens using GP+2-Fer and GP+3-Fer in comparison with GP-Fer.

To test a different adjuvant system, we immunized three groups of mice with GP-Fer, GP+2-Fer or GP+3-Fer adjuvanted with Alhydrogel (alum) and CpG oligodeoxynucleotides (Fig. 4a). Similar to our previous observation with MPLA/Quil-A (**Fig. S4a**), GP-Fer elicited higher EBOV GP-specific IgG titers than GP+2-Fer or GP+3-Fer at early time points (**Fig. S5a**), while mice from all groups developed similar responses after two boosts (Fig. 4b**, Fig. S5b,c**). Unlike the previous immunization with MPLA/Quil-A, where mouse antisera neutralized EBOV one week after the first boost (Fig. 3f), there was a delay in developing strong neutralizing activity with alum/CpG, which was measurable three weeks after the first boost (Fig. 4c). Nonetheless, antisera from all mice potently neutralized EBOV after two booster injections on weeks 3 and 6 (**Fig. S5d**). Despite cross-binding of BDBV GP and SUDV GP (Fig. 4d), only six or two of nine mice in the GP-Fer group cross-neutralized BDBV or SUDV, respectively (Fig. 4e**, Fig. S5e,f**). By contrast, all mice in the GP+2-Fer and the GP+3-Fer groups showed cross-neutralizing activity against BDBV and SUDV. These results again confirmed that a three-dose regimen with hyperglycosylated GP-Fer could induce cross-neutralizing antisera regardless of the different adjuvants used.

**Figure 4.**
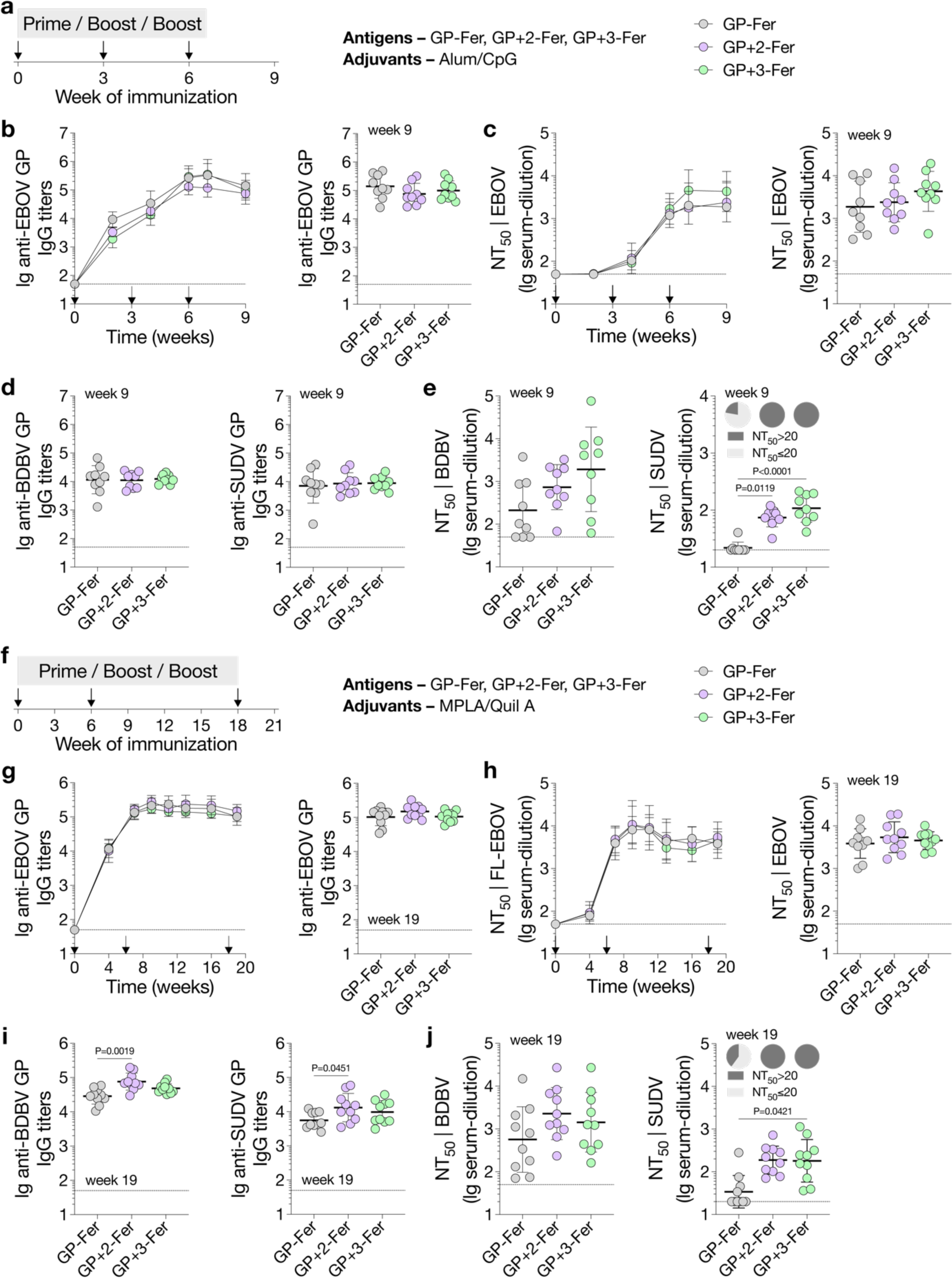
Cross-neutralizing responses elicited by hyperglycosylated GP-Fer under different immunization regimens. **a**, Mouse immunization with GP-Fer and its glycan variants (GP+2-Fer or GP+3-Fer) with a three-dose regimen on days 0, 21 and 42. Mice were immunized with 5 µg of antigens adjuvanted with alum/CpG (500 µg/20 µg) *via* subcutaneous injections (*n*=9 per group). **b**, EBOV GP-specific IgG titers over time (left). EBOV GP-specific IgG titers of individual mice on day 63 (week 9, right). **c**, NT_50_ against EBOV over time (left). NT_50_ against EBOV from individual mice on day 63 (right). **d**, BDBV GP-(left) or SUDV GP-specific (right) IgG titers of individual mice on day 63. **e**, NT_50_ against BDBV (left) or SUDV (right) from individual mice on day 63. Pie charts indicate the percentage of mice that developed neutralizing activity (NT_50_ over 20-fold serum dilution) against SUDV. **f**, Mouse immunization with GP-Fer and its glycan variants (GP+2-Fer, GP+3-Fer or GP+4-Fer) with a three-dose regimen on days 0, 42 and 126. Mice were immunized with 5 µg of antigens adjuvanted with MPLA/Quil-A (10 µg/10 µg) *via* subcutaneous injections (*n*=10 per group). **g**, EBOV GP-specific IgG titers over time (left). EBOV GP-specific IgG titers of individual mice on day 133 (week 19, right). **h**, NT_50_ against EBOV over time (left). NT_50_ against EBOV from individual mice on day 133 (right). **i**, BDBV GP-(left) or SUDV GP-specific (right) IgG titers of individual mice on day 133. **j**, NT_50_ against BDBV (left) or SUDV (right) from individual mice on day 133. Pie charts indicate the percentage of mice that developed neutralizing activity (NT_50_ over 20-fold serum dilution) against SUDV. Each circle represents a single mouse. Horizontal dashed lines indicate the limits of quantification. Data are presented as geometric mean ± standard deviation of log-transformed data in **b**-**e** and **g**-**j**. Comparisons of means of the treatment groups and the control group were performed using one-way ANOVA with a Dunnett’s test in **e,i,j**. *P* values of 0.05 or less are considered significant and plotted.

We further investigated a different immunization regimen with delayed boosts(62, 63) in contrast to the two immunization regimens above, which involved three injections within six weeks. We immunized three groups of mice with GP-Fer, GP+2-Fer or GP+3-Fer adjuvanted with MPLA/Quil-A on weeks 0, 6 and 18 (**Fig. 4f**), since this adjuvant system induced more consistent responses within each group than alum/CpG did. Despite similar titers against EBOV GP at week four (**Fig. S6a**), GP+2-Fer and GP+3-Fer groups induced significantly higher cross-binding titers against BDBV GP and SUDV GP (**Fig. S6b,c**). The second injection at week six elevated EBOV GP-specific IgG titers (**Fig. 4g**) and substantially increased the neutralizing response against EBOV in all groups (**Fig. 4h, Fig. S6d**). This delayed boost also induced cross-neutralizing activity against BDBV and SUDV, particularly for GP+2-Fer and GP+3-Fer (**Fig. S6e,f**). Antibody titers and NT_50_ remained at high levels with negligible differences among groups for more than ten weeks before a second boost. Similar to previous results (**Fig. 3h**, **Fig. 4d**), mice from all groups showed cross-binding titers toward BDBV GP and SUDV GP at one week post-second boost (**Fig. 4i**). Despite a minimal effect on NT_50_ against EBOV (**Fig. S6g**), all mice developed cross-neutralizing activity against BDBV with the lowest titers in the GP-Fer group (**Fig. 4j, Fig. S6h**). In addition, all mice in the GP+2-Fer and GP+3-Fer groups cross-neutralized SUDV, whereas only four of ten mice in the GP-Fer group did (**Fig. S6i**). These immunization studies further validated that hyperglycosylated GP-Fer (GP+2-Fer and GP+3-Fer) induced superior cross-neutralizing activity than GP-Fer did.

## DISCUSSION

Ebola virus outbreaks emerge periodically in Africa and cause severe morbidity and high mortality in infected individuals(1–3). Despite the successful development of two viral vaccines(4), they are only distributed to high-risk groups during active outbreaks(8, 9), partially due to challenges of cold-chain storage and transport(5). In addition, the two vaccines are only approved to protect against EBOV(10, 11), one of three *orthoebolavirus* species with epidemic and pandemic potential(2, 3). Indeed, recent outbreaks of SUDV with high case-fatality rates highlight our vulnerability, where no licensed vaccines or therapeutics were available(64, 65), although viral vaccine candidates have shown promise in preclinical studies(66). Therefore, there is an urgent need to develop effective vaccines that go beyond species-specific protection and confer cross-protective immunity against different *orthoebolavirus* species.

Because *orthoebolavirus* species are antigenically distinct, current vaccine candidates that encode individual GP sequences primarily elicit species-specific responses with little cross-protection(66). As a result, heterologous vaccines that contain viral particles presenting different GPs have been used to induce cross-protection preclinically(41–44). Nonetheless, these multivalent viral vaccines still require cold-chain storage, complicated manufacturing processes and vaccination regimens that are challenging to implement in remote areas(5). Even for the monovalent viral vaccine (Ervebo), its storage and transport require temperatures below −60 °C(5), which poses significant logistical challenges in African countries with a high year-round ambient temperature and a high risk of Ebola virus outbreaks. By contrast, protein-based immunogens represent an important alternative for widespread use. We recently developed a ferritin-based COVID-19 vaccine that maintains potency for at least two weeks at 37°C(63). This vaccine also induced potent, durable and broad antibody responses against existing variants of concern and SARS-CoV. Therefore, GP-based immunogens that share similar thermostability and elicit cross-protective responses hold promise as effective universal *orthoebolavirus* vaccine candidates.

In this work, we harnessed hyperglycosylation as an immunofocusing approach to design epitope-focused antigens based on EBOV GP, given that previous studies on influenza(47, 67) and HIV-1(45, 68) have demonstrated success of hyperglycosylation in focusing humoral immune responses toward particular epitopes of interest. We first determined potential glycosylation sites on GP and then created three hyperglycosylated GP variants with different levels of epitope masking of the poorly conserved glycan cap region. To increase immunogenicity, we displayed these GP variants on Fer for subsequent immunization studies. With a prime-boost regimen, we found that GP-Fer induced substantially higher neutralizing antibody responses than GP trimer did, although a single dose of GP trimer was previously shown to induce a similar level of antibody responses to GP-Fer over time(69). Compared with wild-type GP-Fer that induced minimal cross-neutralizing activity, both GP+2-Fer and GP+3-Fer consistently elicited cross-neutralizing antisera against BDBV and SUDV with different adjuvants and immunization regimens. Although GP+4 was most effective at shielding antibodies that target the glycan cap (*e.g.*, REGN3470), it induced the weakest cross-neutralizing activity among all groups, possibly due to over-masking of nearby epitopes or decreased thermostability.

We observed weaker cross-neutralizing activity against SUDV than BDBV, which correlates with the sequence conservation of GP – SUDV GP is more divergent from EBOV GP in its protein sequence than BDBV GP. In a screen for cross-reactive antibodies from an Ebola virus disease survivor library, 72% of the antibodies recognized BDBV GP, while only 11% cross-reacted with SUDV GP(27). In addition, although antibody-dependent enhancement is a theoretical concern over Ebola vaccine development and has been reported *in vitro* with weakly or non-neutralizing antibodies(70) or antisera(35), we did not observe this effect in any of our immunization studies against the three *orthoebolavirus* species we tested.

Besides neutralizing activity, total antibody-binding levels are regarded as a strong correlate of protection, considering Fc-mediated effector functions as another important route to protection(71–73). Whereas all antigens studied here induced high titers of cross-reactive antibodies that bound BDBV and SUDV GP, the cross-neutralizing activity in antisera raised against GP+2-Fer and GP+3-Fer provides a potentially direct mechanism of protection against viral infection. It will be important to investigate the mechanism of protection by our vaccine candidates in future heterologous lethal challenge studies.

## ACKNOWLEDGMENTS

We thank members of the Kim Lab for fruitful discussions on the project. We thank Drs. Mirella Bucci, Soohyun Kim, Theodora U.J. Bruun, Thuy-Tien Thi Nguyen and Hyeonseob Lim for comments on the manuscript. D.X. and A.E.P. thank Nicholas M. Riley for helping with glycosylation analysis. D.X. thanks Zheng Cao for technical assistance with transmission electron microscopy and dynamic light scattering. D.X. and A.E.P. acknowledge the postdoctoral fellowship from the Stanford Maternal and Child Health Research Institute. A.U. acknowledges the support from Stanford Medical Scientist Training Program (T32 GM007365). J.J.P and R.A.D were supported by an NIH grant (UC7 AI095321). J.I.M. and N.J.S. acknowledge support from the Boston University and the Edward Avedisian Professorship. This work was supported by an NIH Director’s Pioneer Award (DP1-AI158125), the Virginia & D.K. Ludwig Fund for Cancer Research, and the Chan Zuckerberg Biohub to P.S.K.

## AUTHOR CONTRIBUTIONS

Designed research: D.X., A.E.P., P.S.K.

Performed research: D.X., A.E.P., A.U., M.S., J.D., J.J.P.

Analyzed data: D.X., A.E.P., J.J.P., R.A.D., J.I.M., N.J.S., P.S.K.

Wrote the paper: D.X., P.S.K.

## COMPETING INTERESTS

D.X., A.E.P. and P.S.K. are named as inventors on a patent application applied for by Stanford University and the Chan Zuckerberg Biohub on immunogenic ebolavirus fusion proteins and related methods. A.E.P. is an employee of, and P.S.K. is a co-founder and member of the Board of Directors of Vaccine Company, Inc. N.J.S. is a named inventor for filovirus vaccines. All other authors declare no competing interests.

## MATERIALS AND METHODS

### Cell lines

HEK-293T and Vero E6 cells were purchased from American type culture collection (ATCC) and maintained in D10 medium – Dulbecco’s Modified Eagle Medium (DMEM, Cytiva) supplemented with 10% fetal bovine serum (GeminiBio) and 1% *L*-glutamine/penicillin/streptomycin (GeminiBio). Expi-293F cells were maintained in Freestyle293/Expi-293 media (2:1, *v/v*, Thermo Fisher) in polycarbonate shaking flasks (Triforest Labware).

### Antibodies

Monoclonal antibodies against EBOV GP (REGN3471, FVM04, REGN3470, ADI-15742, KZ52, and ADI-16061) were expressed in Expi-293F cells *via* transient transfection. A mouse anti-EBOV GP antibody (IBT BioServices, 0201-020) was used for staining in serum neutralization assays with wild-type EBOV. Goat anti-mouse IgG, HRP conjugated (BioLegend, 405306) or rabbit anti-human IgG, HRP conjugated (Abcam, ab6759) were used as secondary antibodies for Western blots or enzyme-linked immunosorbent assays (ELISAs).

### Antigen and antibody cloning

DNA encoding the EBOV GP (Mayinga, 1976) ectodomain with the mucin-like domain deleted (residues 1-308, 490-656) and the transmembrane domain replaced with a GCN4(74) or foldon trimerization(75) domain followed by an Avi-Tag(76) and a hexahistidine tag was cloned into a mammalian protein expression vector (pADD2) by In-Fusion (Takara Bio).

Similarly, DNA encoding SUDV GP (Uganda/Gulu, 2000) ectodomain with the mucin-like domain deleted (residues 1-345, 506-656) or BDBV GP (Uganda/Butalya, 2007) ectodomain with the mucin-like domain deleted (residues 1-312, 471-640) was cloned into pADD2 vector with a GCN4 or foldon trimerization domain followed by an Avi-Tag and a hexahistidine tag on the C-terminus. GP-Fer was constructed by In-Fusion cloning of EBOV GP (residues 1-308, 491-656) and *H. pylori* ferritin(49) (residues 5-168) with a Ser-Gly-Gly linker into the pADD2 vector. GP-Fer variants were constructed by site-directed mutagenesis using GP-Fer as the backbone. DNA fragments encoding the variable heavy chain (HC) and light chain (LC) were codon-optimized and synthesized by Integrated DNA Technologies. Fragments were inserted into an expression plasmid containing VRC01 HC and LC constant domains by In-Fusion.

All plasmids were sequence-confirmed by Sanger sequencing (Sequetech). For transfection purposes, plasmids were transformed into Stellar cells (Takara Bio), isolated by Maxiprep kits (Macherey Nagel), filtered through a sterile 0.45-µm membrane in a biosafety cabinet, and stored at −20°C.

### Protein expression and purification

All antigens and antibodies were expressed in Expi-293F cells. Expi-293F cells were cultured at 37°C under constant shaking (120 rpm) in a humidified CO_2_ (8%) incubator. Expi-293F cells were transfected at a density of 3-4 × 10^6^ cells/mL. For 200 mL transfection, the transfection mixture was made by adding 120 µg plasmid DNA (from Maxiprep) into 20 mL expression media, followed by the dropwise addition of 260 µL FectoPro transfection reagent (Polyplus) with vigorous mixing. For antibody production in 200 mL Expi-293F cells, the transfection mixture contained 60 µg light-chain plasmid DNA and 60 µg heavy-chain plasmid DNA. Transfection mixtures were incubated at room temperature for 10 min before being transferred to Expi-293F cells. *D*-glucose (4 g/L, Sigma-Aldrich) and valproic acid (3 mM, Acros Organics) were added to the cells immediately post-transfection to increase recombinant protein production. Cells were boosted again with *D*-glucose three days post-transfection and harvested on day four by centrifugation at 7000 ×*g* for five min. The supernatant was filtered through a 0.22-µm membrane for subsequent purification processes. Biotinylated GP was expressed using the same protocol but in the presence of the BirA enzyme.

His-tagged proteins were purified with HisPur^TM^ Ni-NTA resin (Thermo Fisher). Briefly, filtered supernatant from Expi-293F cells was mixed with Ni-NTA resin (1 mL resin per liter supernatant) and incubated at 4°C overnight. The mixture was then passed through a gravity-flow column, washed with 20 mM imidazole in HEPES buffer saline (HBS, 20 mM HEPES, pH 7.4, 150 mM NaCl), and then eluted with 250 mM imidazole in HBS. Elution was concentrated with centrifugal filters (50 kDa MWCO, Millipore Sigma) and buffer-exchanged into HBS for size-exclusion chromatography using a Superose^®^ 6 Increase 10/300 GL column (Cytiva). Peak fractions were pooled, concentrated, buffer-exchanged to HBS with 10% glycerol, and filtered through a 0.22-µm membrane.

GP-Fer and its glycan variants were purified by anion-exchange chromatography followed by size-exclusion chromatography. Filtered supernatant from Expi-293F cells was directly applied to a HiTrap Q HP column (Cytiva) on an ÄKTA Protein Purification System (Cytiva). Columns were washed with Tris buffer (20 mM, pH 8.0), and protein nanoparticles were eluted with a NaCl gradient (0 - 1.0 M). Nanoparticle-containing fractions were determined by Western blotting with mAb114, pooled and concentrated using centrifugal filters (100 kDa MWCO), and subsequently purified twice with size exclusion chromatography using an SRT SEC-1000 column (Sepax Technologies). Peak fractions were pooled, concentrated, buffer-exchanged to HBS with 10% glycerol, and filtered through a 0.22-µm membrane.

All antibodies were purified with MabSelect PrismA protein A chromatography. Filtered supernatant from Expi-293F cells was directly applied to a MabSelect PrismA column (Cytiva) on an ÄKTA Protein Purification System. Columns were washed with HBS, and then antibodies were eluted with glycine (100 mM, pH 2.8) into HEPES buffer (1 M, pH 7.4). Fractions were concentrated and buffer-exchanged to HBS with 10% glycerol.

The concentration of all proteins was determined by absorbance at 280 nm, and the purity was assessed by protein gel electrophoresis. Protein samples were flash-frozen in liquid nitrogen and stored at −20°C.

### Glycan screening

Based on the structure of EBOV GP (PDB ID: 5JQ3), we selected 16 sites on the surface of each GP protomer for potential glycosylation, where an *N*-linked glycosylation site (N-X-S/T, where X can be any amino acid except proline) was introduced by site-directed mutagenesis. We then compared the expression level of all glycan variants to wild-type GP in Expi-293F cells *via* immunoblotting. Three days after transient transfection, Expi-293F cells were harvested *via* centrifugation at 7,000 ×*g* for 5 min. Supernatant samples were collected, mixed with Laemmli loading buffer (4×, Bio-Rad), boiled at 95°C for 5 min, and then loaded onto protein gels (4-20% Mini-PROTEAN^®^ TGX^TM^ Precast Gels, Bio-Rad). After gel electrophoresis, proteins were transferred to 0.2-µm nitrocellulose membranes (Bio-Rad), and these blots were blocked with PBST (phosphate-buffered saline, 0.1% Tween-20) with milk (10% *w*/*v*, nonfat dry milk, Bio-Rad) for one hour at room temperature. Primary antibody – mAb114 (2 mg/mL, 1:4000 dilution in PBST with milk) was then added for one-hour incubation, followed by rabbit anti-human IgG, HRP conjugated (1:4000 dilution in PBST with milk) as the secondary antibody (45-min incubation). Blots were rinsed in PBST for five min between steps and finally developed using a Western blotting substrate (Pierce ECL, Thermo Scientific). Imaging was acquired on a chemiluminescence imager (GE Amersham Imager 600) and analyzed with Fiji (ImageJ v2.3.0). Glycan variants that retained >50% of GP expression were considered “good glycans”.

### Consurf analysis

Protein sequences of GP from six *orthoebolavirus* species (Ebola – AAG40168.1, Sudan – AAU43887.1, Bundibugyo – AYI50307.1, Täi Forest – AAB37093.1, Reston – AAC54891.1 and Bombali – ASJ82195.1) were aligned using Clustal Omega to create a multiple sequence alignment (MSA). The MSA was analyzed by the Consurf server(77) (https://consurf.tau.ac.il/) based on the structure of EBOV GP (PDB ID: 5JQ3) and residues in the modified PDB file were re-colored in PyMOL (Schrödinger) using a python script. A nine-color histogram was used to evenly assign a range of conservation scores to a range of colors (blue – conserved, red – variable).

### Differential scanning fluorimetry

Thermal melting profiles of proteins were measured by differential scanning fluorimetry on a Prometheus NT.48 instrument (NanoTemper). Protein samples (0.1 mg/mL) were loaded into glass capillaries (NanoTemper) and then subjected to a temperature gradient from 20 to 95°C at a heating rate of 1°C per min. Intrinsic fluorescence (350 nm and 330 nm) was recorded as a function of temperature. Thermal melting curves were plotted using the first derivative of the ratio (350 nm/330 nm). Melting temperatures were calculated automatically by the instrument (PR.ThermControl software) and represented peaks in the thermal melting curves.

### Antibody ELISAs

Nunc 96-well MaxiSorp plates (Thermo Fisher) were coated with streptavidin (4 µg/mL in DPBS, 60 µL per well, Thermo Fisher) for one hour at room temperature. Plates were washed three times with Milli-Q H_2_O (300 µL) using a plate washer (ELx405 BioTek) and then blocked with ChonBlock (120 µL per well, Chondrex) overnight at 4°C. For subsequent steps, all dilutions were made in DPBS with 0.05% Tween-20 and 0.1% BSA, and ELISA plates were rinsed with PBST (300 µL, three times) in between. Biotinylated EBOV GP proteins (wild-type, GP+2, GP+3, or GP+4 at 2 µg/mL) were added to the plates and incubated for one hour at room temperature. Then, serially diluted monoclonal antibodies (mAbs, starting from 200 nM, followed by 10-fold dilution) were added to the plates and incubated for another hour. Rabbit anti-human IgG, HRP-conjugated (1:4,000) was added for one-hour incubation before rinsing with PBST six times. ELISA plates were developed with the TMB substrate (1-Step Turbo-TMB, Thermo Fisher) for six minutes and terminated with sulfuric acid (2M). Absorbance at 450 nm (A450) was recorded on a microplate reader (Synergy^TM^ HT, BioTek).

To test antibody binding to GP-Fer and its glycan variants, these protein nanoparticles (2 µg/mL in DPBS, 60 µL per well) were directly coated onto Nunc 96-well MaxiSorp plates for one hour at room temperature. Plates were washed three times with Milli-Q H_2_O (300 µL) and then blocked with ChonBlock (120 µL per well) overnight at 4°C. Serially diluted mAbs were added to the plates, followed by secondary antibodies, TMB substrates, and sulfuric acid. Absorbance at 450 nm (A450) was recorded on a microplate reader (Synergy^TM^ HT, BioTek).

### Size-exclusion chromatography–multi-angle light scattering (SEC-MALS) analysis

SECMALS analysis of protein nanoparticles was performed on a 1260 Infinity II high-performance liquid chromatography system (Agilent) coupled with a miniDAWN and Optilab detectors (Wyatt Technologies) for light scattering and refractive index analysis. Purified GP-Fer, its glycan variants, and wild-type ferritin (10 µg of each sample) were loaded onto an SRT SEC-1000 column (4.6 × 300 mm, Sepax) sequentially for analysis. ASTRA software (Wyatt Technologies) was used for quantitative analysis of the molar mass of protein nanoparticles.

### Dynamic light scattering and Transmission electron microscopy

Protein nanoparticles (GP-Fer, GP+2-Fer, GP+3-Fer, or GP+4-Fer) were diluted to 0.05 mg/mL with DPBS and filtered through a 0.22-µm membrane before analysis. The hydrodynamic size of protein nanoparticles was then measured on a Zetasizer Nano ZS instrument with a 10-mW helium-neon laser and thermoelectric temperature controller (Malvern Panalytical). Before each sample, the temperature of the instrument was equilibrated for one min at 25 °C.

Protein nanoparticles (0.1 mg/mL, 10 µL) were pipetted onto a carbon-coated copper grid (Ted Pella) and incubated for five min, followed by negative staining with uranyl acetate (2%, *w/v*) for 1.5 min. TEM grids were dried overnight, and images were acquired on a Tecnai T12 cryo-electron microscope (FEI) operating with an acceleration voltage of 120 kV.

### Mouse immunization studies

All animals were maintained in accordance with the Public Health Service Policy for “Human Care and Use of Laboratory Animals” under a protocol approved by the Stanford University Administrative Panel on Laboratory Animal Care (APLAC-33709). Female BALB/c mice (6-8 weeks) were purchased from Jackson Laboratories, and female C57BL/6 mice (6-8 weeks) were purchased from Jackson Laboratories or Charles River. To compare GP to GP-Fer, we immunized two groups of BALB/c mice (*n*=10) with 5 µg protein antigens adjuvanted with 10 µg Monophosphoryl lipid A (MPLA, Invivogen) and 10 µg Quil-A (Invivogen) *via* subcutaneous injection on day 0 and 21. To compare GP-Fer variants, we immunized four groups of C57BL/6 mice (*n*=10) with 5 µg protein antigens adjuvanted with 10 µg MPLA and 10 µg Quil-A *via* subcutaneous injection on day 0, 21, and 42. To examine a different set of adjuvants, we immunized three groups of C57BL/6 mice (*n*=9) with 5 µg protein antigens adjuvanted with 500 µg alum (Alhydrogel^®^, Invivogen) and 20 µg CpG (ODN 1826, Invivogen) *via* subcutaneous injection on day 0, 21, and 42. To investigate a different immunization regimen, we immunized three groups of C57BL/6 mice (*n*=10) with 5 µg protein antigens adjuvanted with 10 µg MPLA and 10 µg Quil-A *via* subcutaneous injection on day 0, 42, and 126. Pre-immune, interim and final blood samples were collected by retro-orbital bleeding into serum gel tubes (Sarstedt). Serum gel tubes were centrifuged at 10,000 ×*g* for 6 min, and sera were collected and stored at −80 °C.

### Serum ELISAs

Nunc 96-well MaxiSorp plates were coated with streptavidin (4 µg/mL in DPBS, 60 µL per well) for one hour at room temperature. These plates were washed three times with MilliQ-H_2_O (300 µL) using a plate washer and then blocked with ChonBlock (120 µL per well) overnight at 4°C. For subsequent steps, all dilutions were made in DPBS with 0.05% Tween-20 and 0.1% BSA, and ELISA plates were rinsed with PBST (300 µL, three times) in between.

Biotinylated EBOV GP (wild-type, 2 µg/mL) was added to the plates and incubated for one hour at room temperature. To test cross-reactive binding, we used biotinylated SUDV or BDBV GP (2 µg/mL) instead of EBOV GP for this step. Mouse antisera were serially diluted (5-fold dilution) and then added to the ELISA plates for one-hour incubation at room temperature. Goat anti-mouse IgG, HRP-conjugated (1:4,000) was added for one-hour incubation before rinsing with PBST six times. ELISA plates were developed with the TMB substrate for six minutes and terminated with sulfuric acid (2M). Absorbance at 450 nm (A450) was recorded on a microplate reader (Synergy^TM^ HT, BioTek).

### Production of pseudotyped lentiviruses

EBOV and SUDV GP-pseudotyped lentiviruses encoding a luciferase-ZsGreen reporter were produced in HEK-293T cells by co-transfection of five plasmids(78). This five-plasmid system includes a packaging vector (pHAGE-Luc2-IRES-ZsGreen, a plasmid encoding full-length EBOV or SUDV GP (pCDNA3.1 EBOV FL-GP or pCDNA3.1 SUDV FL-GP), and three helper plasmids (pHDM-Hgpm2, pHDM-Tat1b, and pRC-CMV_Rev1b). One day before transfection, HEK-293T cells were seeded in 10-cm Petri dishes (5 × 10^6^ cells per Petri dish). Transfection mixture was prepared by adding five plasmids (10 µg packaging vector, 3.4 µg GP-encoding plasmid, and 2.2 µg of each helper plasmid) to 1 mL D10 medium, followed by the addition of BioT transfection reagent (30 µL, Bioland Scientific) in a dropwise manner with vigorous mixing. After 10-min incubation at room temperature, the transfection mixture was transferred to HEK-293T cells in the Petri dish. Culture medium was replenished 24 hours post-transfection, and after another 48 hours, viruses were harvested and filtered through a 0.45-µm membrane.

BDBV GP-pseudotyped lentiviruses were produced using the same method but in Expi-293F cells. Expi-293F cells were adjusted to 3 × 10^6^ cells per mL one day before transfection. For 200 mL Expi-293F cells, transfection mixture was prepared by adding five plasmids (200 µg packaging vector, 68 µg plasmid encoding BDBV GP with the mucin domain deleted – Δ313-470, and 44 µg of each helper plasmid) to 20 mL FreeStyle293/Expi-293 media, followed by the addition of 600 µL BioT transfection reagent in a dropwise manner with vigorous mixing. After 10-min incubation at room temperature, the transfection mixture was transferred to Expi-293F cells. Expi-293F cells were immediately boosted with *D*-glucose (4 g/L) and valproic acid (3 mM). After 72 hours, viruses were harvested and filtered through a 0.45-µm membrane.

All pseudotyped lentiviruses (EBOV, SUDV, or BDBV) were aliquoted, flash-frozen in liquid nitrogen, stored at −80°C, and titrated in HEK-293T cells before further use.

### Pseudoviral neutralization assays with monoclonal antibodies

Neutralization of three pseudotyped lentiviruses (EBOV, SUDV, or BDBV) was validated with monoclonal antibodies (mAbs) in HEK-293T cells. Cells were seeded in white-walled, clear-bottom 96-well plates at a density of 20,000 cells per well one day before the assay (day 0). On day 1, mAbs (2 µM in HBS with 10% glycerol) were filtered with 0.22-μm sterile membranes and diluted with D10 media.

Subsequently, mAbs were serially diluted (10-fold dilution) in D10 media and mixed with lentiviruses (diluted in D10 medium, supplemented with polybrene, 1:1000, *v/v*) for one hour before being transferred to HEK-293T cells. On day 4, medium was removed, and 100 µL of luciferase substrates (BriteLite Plus, Perkin Elmer) were added to each well. Luminescent signals were recorded on a microplate reader (BioTek Synergy™ HT or Tecan M200). Percent infection was normalized to cells only (0% infection) and virus only (100% infection) on each plate. Neutralization assays were performed in biological replicates with technique duplicates.

### Serum neutralization assays against pseudoviruses

Antisera were heat-inactivated (56°C, 30 min) before neutralization assays. Briefly, HEK-293T cells were seeded in white-walled, clear-bottom 96-well plates (20,000 cells per well) one day before the assay (day 0). On day 1, antisera were serially diluted in D10 media and then mixed with EBOV, SUDV, or BDBV (diluted in D10 medium, supplemented with polybrene, 1:1000, *v/v*) for one hour before being transferred to HEK-293T cells. On day 4, medium was removed, and 100 µL of luciferase substrates (BriteLite Plus, Perkin Elmer) were added to each well. Luminescent signals were recorded on a microplate reader (BioTek Synergy™ HT or Tecan M200). Percent infection was normalized to cells only (0% infection) and virus only (100% infection) on each plate. Neutralization titers (NT_50_) were calculated as the serum dilution where a 50% inhibition of infection was observed. Neutralization assays were performed in technical duplicates.

### Serum neutralization assays against wild-type Ebola viruses

All work with infectious Ebola virus was performed at the National Emerging Infectious Diseases Laboratories (NEIDL), Boston University (Boston, MA) in a dedicated BSL-4 biocontainment lab. Vero E6 cells were seeded at a density of 3,500 cells per well into a 384-well plate. The following day, pooled antisera from each group (GP-Fer, GP+2-Fer, GP+3-Fer or GP+4-Fer) and controls were serially diluted in D10, then incubated with wild-type EBOV Mayinga for one hour. The virus-antibody mixture was then transferred to the assay plate and incubated for approximately 20 hours in a humidified CO_2_ (5%) incubator at 37°C. After incubation, the medium was removed and plates were submerged in 10% neutral buffered formalin for inactivation at 4°C for 6 hours. Plates were removed from containment and washed with PBS. Cells were permeabilized with 0.1% Triton X-100 for 15 minutes and blocked with 3.5% BSA for 1 hour at room temperature. Viral protein was stained with a mouse anti-GP antibody (1:2000, IBT 0201-020) for several hours, followed by an Alexa Fluor 488-conjugated, anti-mouse secondary antibody. Nuclei were counterstained with Hoechst 33342. Cells were imaged using an automated microscope (Cytation 1, Agilent-BioTek). Infected and total cells were counted using a customized pipeline in CellProfiler (Broad Institute, available upon request). Percent infection of total cells was calculated and normalized to the untreated controls. Data were plotted with GraphPad Prism.

### Statistical analyses

Statistics were analyzed using GraphPad Prism software (version 9.3.1). Non-transformed data are presented as mean ± standard deviation. Log-transformed data (ELISA titers and NT_50_) are presented as geometric mean ± standard deviation. Comparisons of two groups were performed using the two-tailed Mann–Whitney U test. Comparisons of means of the treatment groups and the control group were performed using one-way ANOVA with a Dunnett’s test. Comparisons of antigen-specific IgG or neutralization titers over time were performed using two-way ANOVA followed by a Bonferroni test. *P* values of 0.05 or less are considered significant and plotted.

### Data Availability

All data supporting the results in this study are available within the main text and its supplementary information. Additional data supporting the results of the study are also available from the corresponding authors upon reasonable request.

## Supplementary Figures

**Figure S1.**
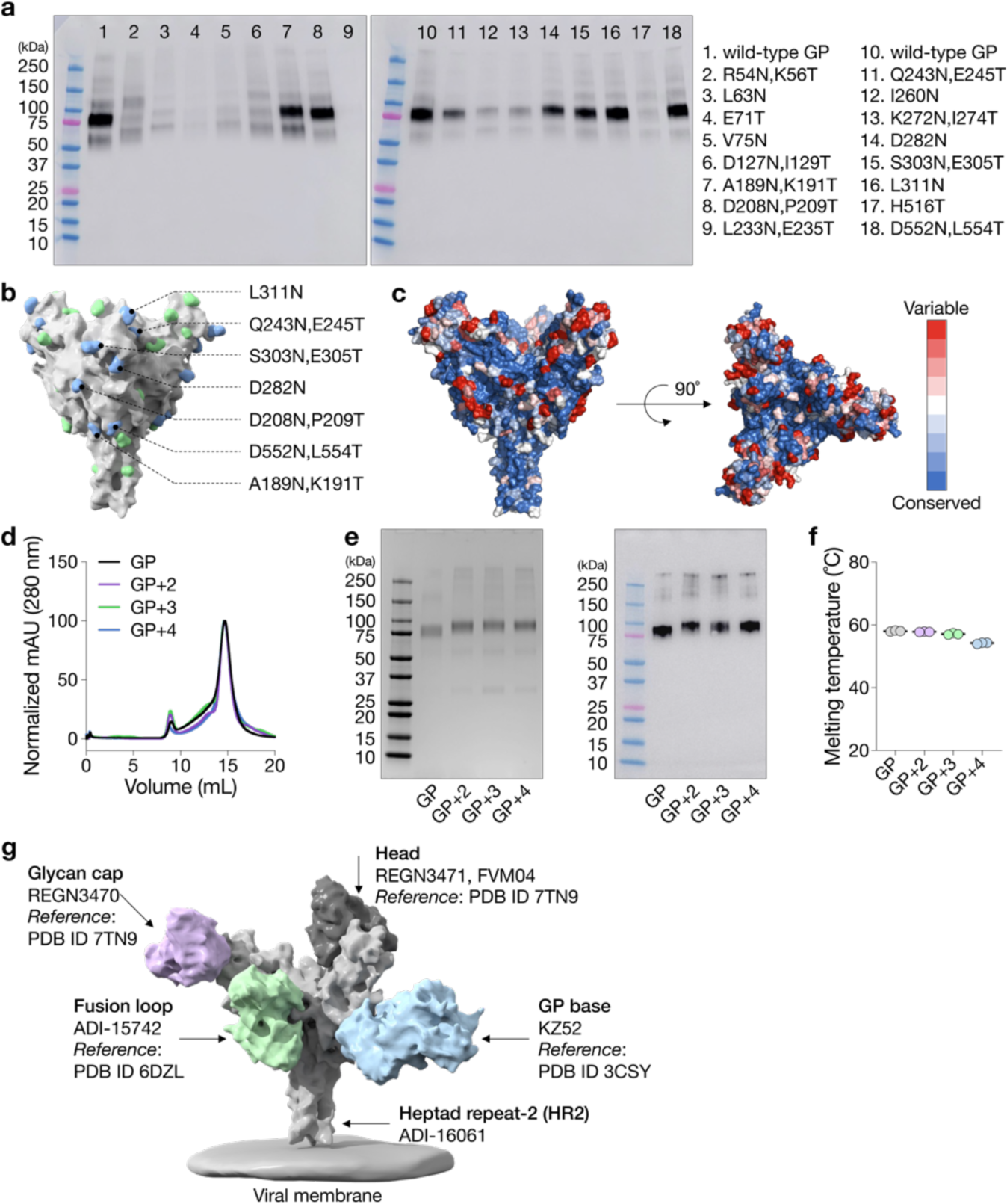
Screening and installation of glycans on EBOV GP. **a**, Analysis of expression levels of single-glycan mutants by Western blots. Supernatant samples from transient transfection were used for the analysis, and blots were detected by mAb114. **b**, Spatial distribution of permissive glycan installations (blue spheres) on the EBOV GP trimer. Green spheres represent endogenous glycans. **c**, The degree of sequence conservation of GP among six *orthoebolavirus* species (Ebola – GenBank: AAG40168.1, Bundibugyo – GenBank: AYI50307.1, Sudan – GenBank: AAU43887.1, Täi Forest – GenBank: AAB37093.1, Reston – GenBank: AAC54891.1 and Bombali – ASJ82195.1). Residues are colored by their conservation scores from the Consurf analysis, where blue and red indicate conserved and variable residues, respectively. **d**, Purification of GP and its glycan variants using size-exclusion chromatography (Superose^®^ 6). Milli-absorbance units (mAU) from the elution profiles of GP and its glycan variants were normalized for comparison. **e**, Protein gel electrophoresis (left) and Western blotting (right) of purified GP and its glycan variants. The protein gel was stained with Coomassie brilliant blue. mAb114 was used as the primary antibody for detection in the Western blot. **f**, Thermal melting temperatures (T_m_) of wild-type GP and its glycan variants measured by differential scanning fluorimetry. Data are presented as mean ± standard deviation (*n*=3 biological replicates). **g**, Different epitopes targeted by the panel of six mAbs used for epitope mapping analysis.

**Figure S2.**
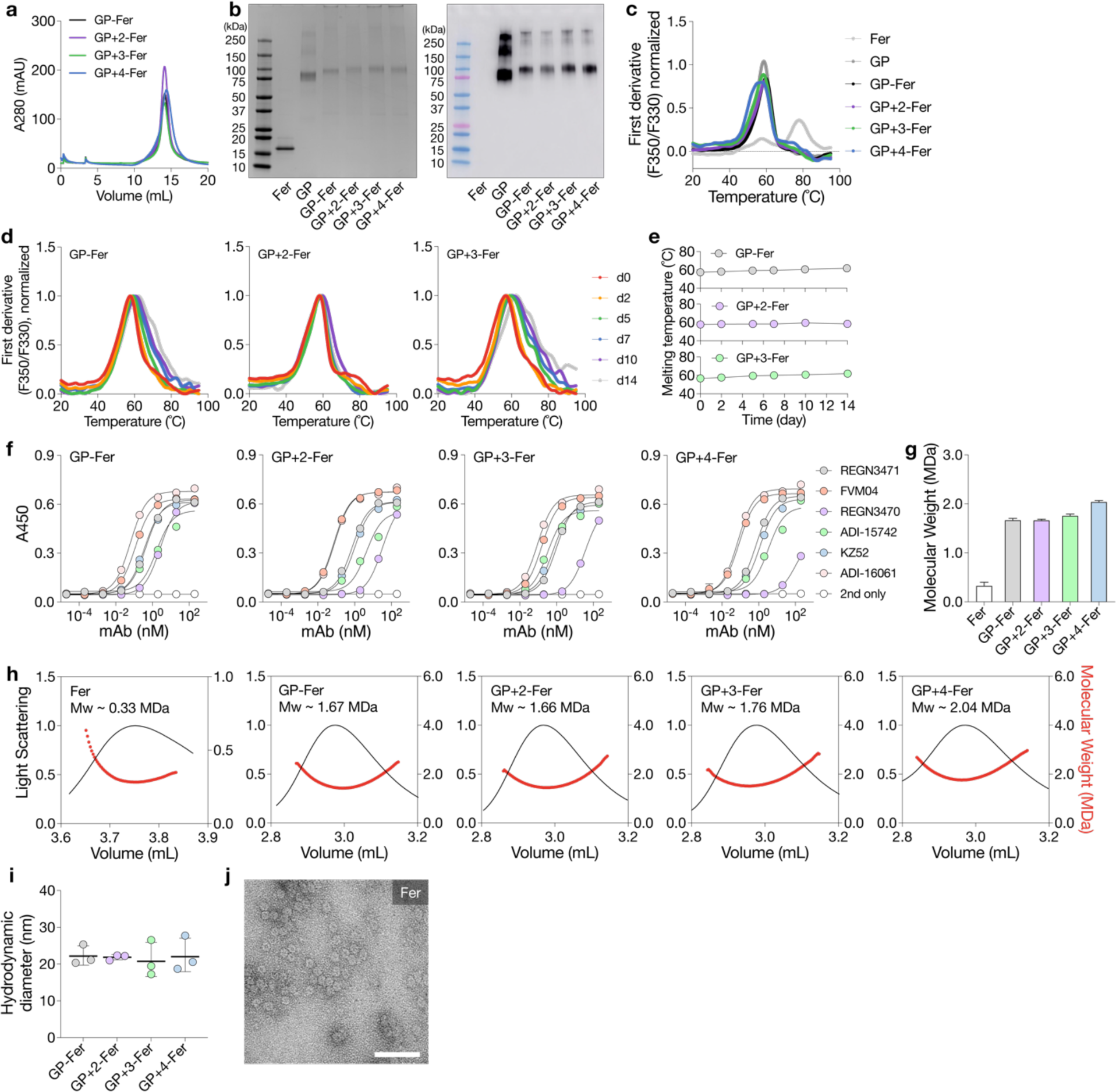
Biochemical and biophysical characterization of GP-Fer and its glycan variants. **a**, Purification of GP and its glycan variants using size-exclusion chromatography (Sepax SRT SEC-1000). **b**, Protein gel electrophoresis (left) and Western blotting (right) of purified GP-Fer and its glycan variants. The protein gel was stained with Coomassie brilliant blue. mAb114 was used as the primary antibody for detection in the Western blot. **c**, Thermal melting profiles of GP-Fer and its glycan variants as measured by differential scanning fluorimetry. Wild-type Fer and GP served as controls. **d,e**, Thermal melting profiles (**d**) and T_m_ (**e**) of GP-Fer, GP+2-Fer and GP+3-Fer after incubation at 37°C for different time (2, 5, 7, 10 or 14 days). **f**, Epitope analysis of GP-Fer and its glycan variants with six GP-specific mAbs by ELISA. Data are presented as mean ± standard deviation (*n*=2 biological replicates with technical duplicates). **g,h**, Molecular weight of Fer, GP-Fer and its glycan variants calculated from the refractive indices and elution profiles by the ASTRA software. **i**, Hydrodynamic diameters of GP-Fer and its glycan variants as measured by dynamic light scattering. Data are presented as mean ± standard deviation (*n*=3 biological replicates). **j**, Transmission electron micrograph of wild-type Fer. Scale bar, 50 nm.

**Figure S3.**
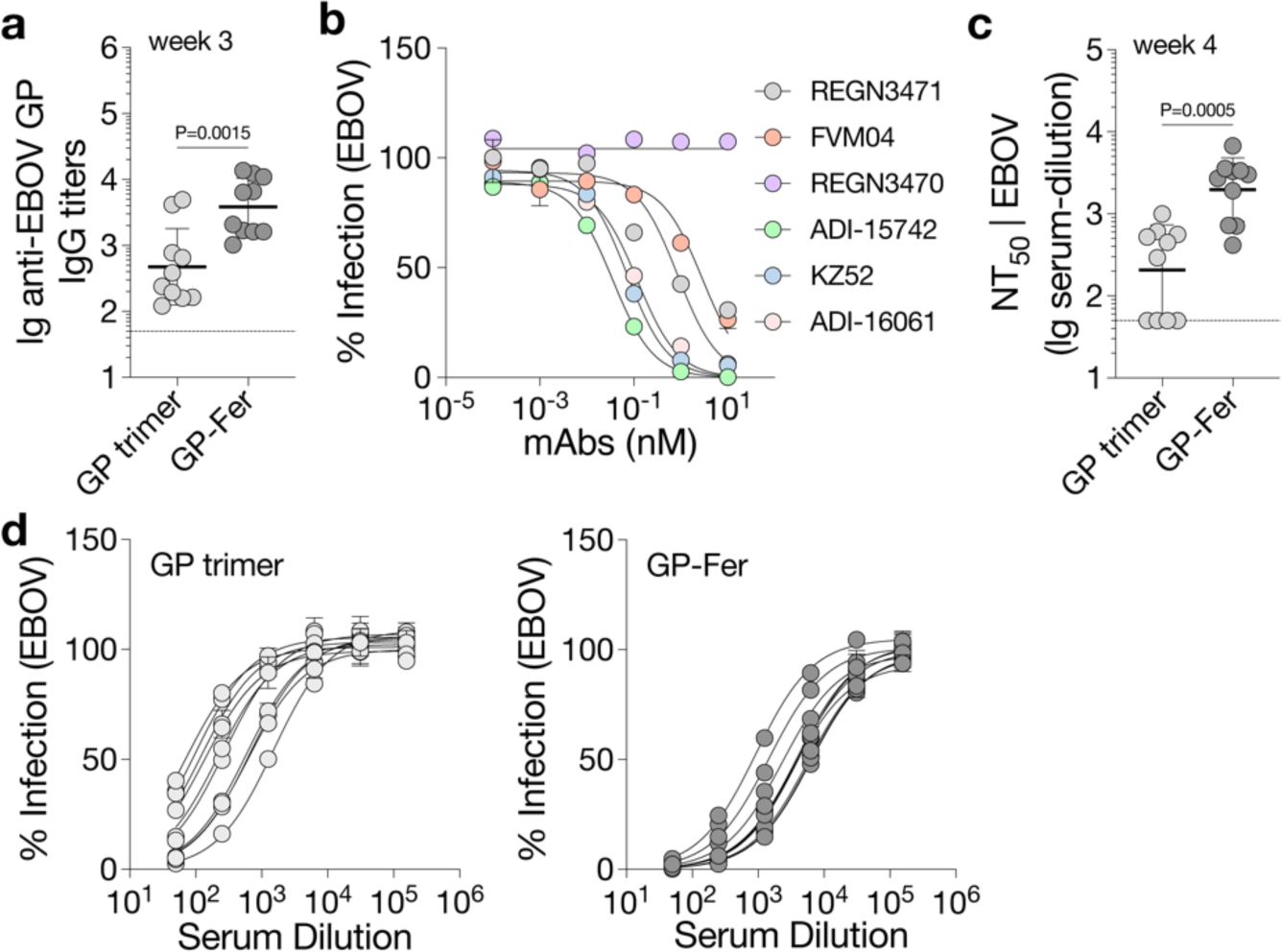
Comparison of GP trimer and GP-Fer in mouse immunization. Mice were immunized as in Fig. 3a with a prime-boost regimen on days 0 and 21 (*n*=10 per group). **a**, EBOV GP-specific IgG titers of individual mice 21 days post-prime. Data are presented as geometric mean ± standard deviation of log-transformed data. Statistical analysis was performed using a two-tailed Mann–Whitney U test. **b**, Validation of neutralization assays against EBOV GP-pseudotyped lentiviruses (EBOV) with six mAbs. REGN3470 is known to be non-neutralizing and serves as a negative control. Data are presented as mean ± standard deviation (*n*=2 biological replicates with technical duplicates). **c**, NT_50_ against EBOV from individual mice on day 28. Data are presented as geometric mean ± standard deviation of log-transformed data. Statistical analysis was performed using a two-tailed Mann-Whitney U test. **d**, Neutralization of EBOV with mouse antisera from the GP trimer (left) or GP-Fer group (right). Each circle or curve represents a single mouse in **a**,**c** and **d**. Dashed lines indicate the limits of quantification. Data are presented as mean ± standard deviation (*n*=2 technical duplicates). *P* values of 0.05 or less are considered significant and plotted.

**Figure S4.**
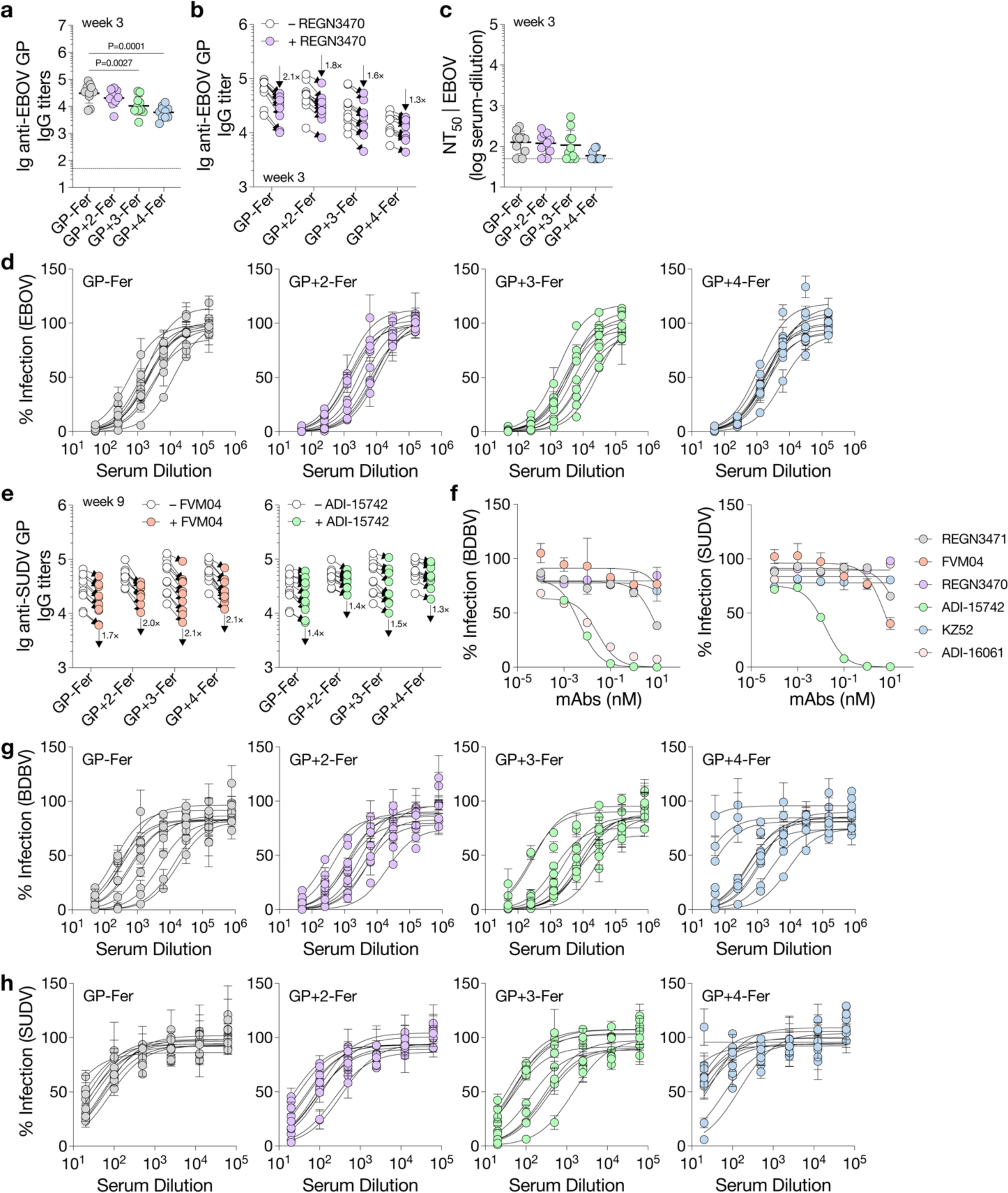
Neutralizing antibody responses elicited by GP-Fer and its glycan variants. Mice were immunized as in Fig. 3d adjuvanted with MPLA/Quil A on days 0, 21 and 42. **a**, EBOV GP-specific IgG titers of individual mice 21 days after the prime. Statistical analysis was performed using one-way ANOVA with a Dunnett’s test. *P* values of 0.05 or less are considered significant and plotted. **b**, Comparison of serum-binding titers to GP in the presence of a competing mAb REGN3470. Arrows indicate overall fold-changes of each group. **c**, NT_50_ against EBOV from individual mice on day 21. **d**, Neutralization of EBOV with mouse antisera from day 63 (*n*=2 technical duplicates). **e**, Comparison of serum-binding titers to GP in the presence of a competing mAb FVM04 (left) or ADI-15742 (right). Arrows indicate overall fold-changes of each group. **f**, Validation of neutralization assays against lentiviruses pseudotyped with BDBV GP (left) or SUDV GP (right) with six mAbs (*n*=2 biological replicates with technical duplicates). **g,h**, Neutralization of BDBV (**g**) or SUDV (**h**) with mouse antisera from day 63 (*n*=2 technical duplicates). Each circle or curve represents a single mouse in **a-e**,**g** and **h**. Dashed lines indicate the limits of quantification. Data are presented as geometric mean ± standard deviation of log-transformed data in **a-c** and **e**. Data are presented as mean ± standard deviation in **d** and **f-h**.

**Figure S5.**
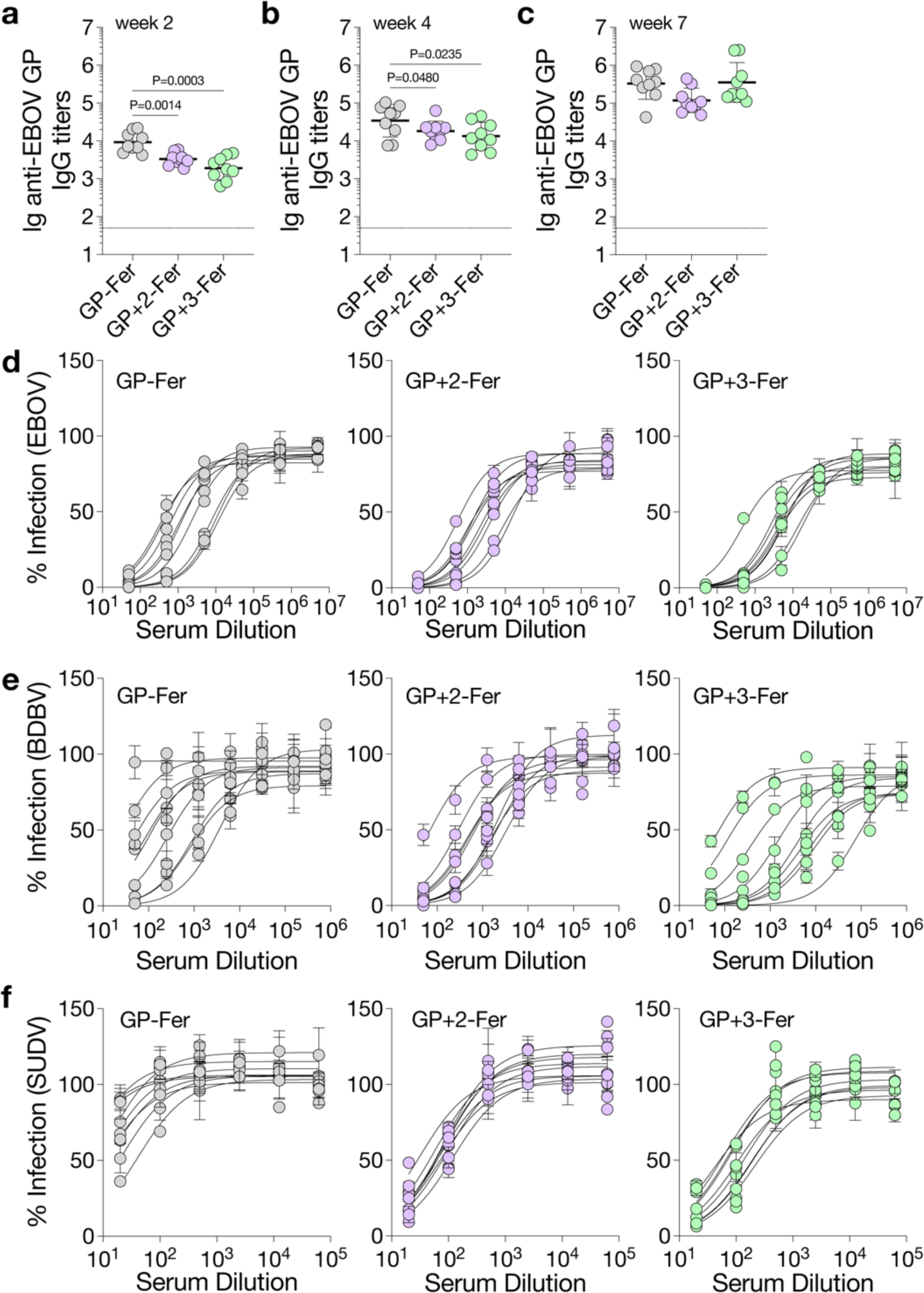
Immunization with GP-Fer and its glycan variants with an alum/CpG adjuvant. Mice were immunized as in Fig. 4a adjuvanted with alum/CpG on days 0, 21 and 42. **a-c**, EBOV GP-specific IgG titers of individual mice on day 14 (**a**), 28 (**b**) or 49 (**c**). Dashed lines indicate the limits of quantification. Data are presented as geometric mean ± standard deviation of log-transformed data. Statistical analysis was performed using one-way ANOVA with a Dunnett’s test. *P* values of 0.05 or less are considered significant and plotted. **d-f**, Neutralization of EBOV (**d**), BDBV (**e**) or SUDV (**f**) with mouse antisera from day 63. Each circle or curve represents a single mouse. Data are presented as mean ± standard deviation (*n*=2 technical duplicates).

**Figure S6.**
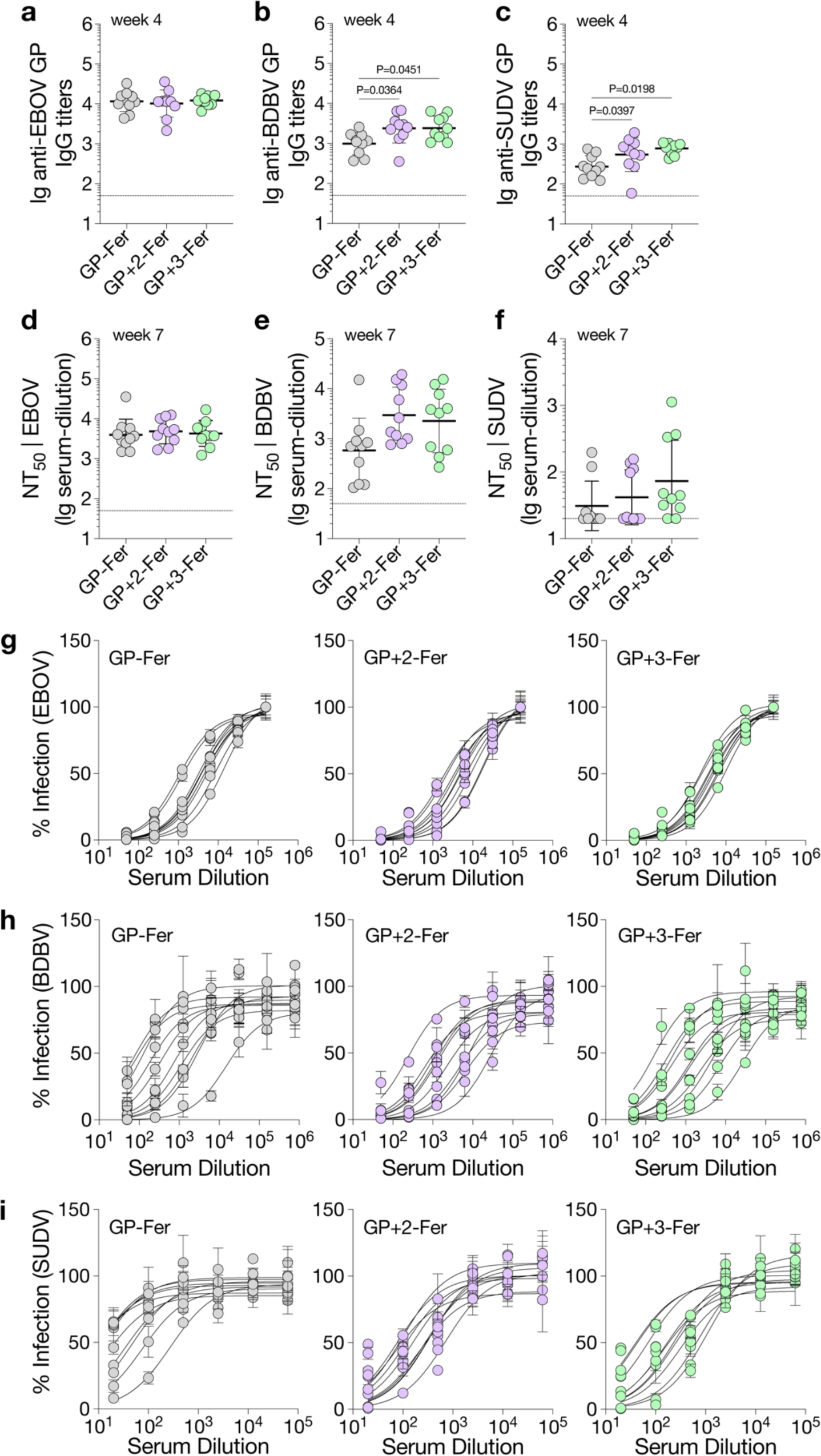
Immunization with GP-Fer and its glycan variants with delayed boosts. Mice were immunized as in Fig. 4f adjuvanted with MPLA/Quil A on days 0, 42 and 126. **a-c**, IgG titers against EBOV GP (**a**), BDBV GP (**b**) or SUDV GP (**c**) of individual mice on day 28. Data are presented as geometric mean ± standard deviation of log-transformed data. Statistical analysis was performed using one-way ANOVA with a Dunnett’s test. *P* values of 0.05 or less are considered significant and plotted. **d-f**, NT_50_ against EBOV (**d**), BDBV (**e**) or SUDV (**f**) from individual mice on day 49. Data are presented as geometric mean ± standard deviation of log-transformed data. **g-i**, Neutralization of EBOV (**g**), BDBV (**h**) or SUDV (**i**) with mouse antisera from day 133. Data are presented as mean ± standard deviation (*n*=2 technical duplicates). Each circle or curve represents a single mouse. Horizontal dashed lines indicate the limits of quantification in **a-f**.

